# Untangling the mechanisms of pulmonary hypertension-induced right ventricular stiffening in a large animal model

**DOI:** 10.1101/2023.04.03.535491

**Authors:** Sotirios Kakaletsis, Marcin Malinowski, Mrudang Mathur, Gabriella P. Sugerman, Jeff J. Lucy, Caleb Snider, Tomasz Jazwiec, Matthew Bersi, Tomasz A. Timek, Manuel K. Rausch

**Affiliations:** Department of Aerospace Engineering & Engineering Mechanics, The University of Texas at Austin, Austin, TX; Division of Cardiothoracic Surgery, Spectrum Health, Grand Rapids, MI; Department of Cardiac Surgery, Medical University of Silesia, Katowice, Poland; Department of Mechanical Engineering, The University of Texas at Austin, TX; Department of Biomedical Engineering, The University of Texas at Austin, TX; Center for Advanced Brain Imaging Research, Rutgers University, New Brunswick, NJ; Department of Mechanical Engineering & Materials Science, Washington University at St. Louis, St. Louis, MO; Department of Cardiac, Vascular and Endovascular Surgery and Transplantology, Medical University of Silesia in Katowice, Silesian Centre for Heart Diseases, Zabrze, Poland

**Author notes:** Corresponding Author: Manuel K. Rausch, PhD, Assistant Professor, The University of Texas at Austin, 2617 Wichita Street, Austin, TX 78712.

## Abstract

**Background:** Pulmonary arterial hypertension (PHT) is a devastating disease with low survival rates. In PHT, chronic pressure overload leads to right ventricle (RV) remodeling and stiffening; thus, impeding diastolic filling and ventricular function. Multiple mechanisms contribute to RV stiffening, including wall thickening, microstructural disorganization, and myocardial stiffening. The relative importance of each mechanism is unclear. Our objective is to use a large animal model as well as imaging, experimental, and computational approaches to untangle these mechanisms.

**Methods:** We induced PHT in eight sheep via pulmonary artery banding. After eight weeks, the hearts underwent anatomic and diffusion tensor MRI to characterize wall thickening and microstructural disorganization. Additionally, myocardial samples underwent histological and gene expression analyses to quantify compositional changes and mechanical testing to quantify myocardial stiffening. All findings were compared to 12 control animals. Finally, we used computational modeling to disentangle the relative importance of each stiffening mechanism.

**Results:** First, we found that the RVs of PHT animals thickened most at the base and the free wall. Additionally, we found that PHT induced excessive collagen synthesis and microstructural disorganization, consistent with increased expression of fibrotic genes. We also found that the myocardium itself stiffened significantly. Importantly, myocardial stiffening correlated significantly with excess collagen synthesis. Finally, our model of normalized RV pressure-volume relationships predicted that myocardial stiffness contributes to RV stiffening significantly more than other mechanisms.

**Conclusions:** In summary, we found that PHT induces wall thickening, microstructural disorganization, and myocardial stiffening. These remodeling mechanisms were both spatially and directionally dependent. Using modeling, we show that myocardial stiffness is the primary contributor to RV stiffening. Thus, myocardial stiffening may be an important predictor for PHT progression. Given the significant correlation between myocardial stiffness and collagen synthesis, collagen-sensitive imaging modalities may be useful for non-invasively estimating myocardial stiffness and predicting PHT outcomes.

## INTRODUCTION

Pulmonary arterial hypertension (PHT) is a devastating disease with a survival rate of only 58% at 3 years^1^. For those eventually succumbing to PHT, complications associated with right heart failure is the predominant cause of death^2, 3^. When failing, the right heart can show hallmarks of both systolic and diastolic dysfunction^4^. Interestingly, recent evidence suggests that measures of diastolic dysfunction may be better predictors for adverse clinical outcomes^5^. That is, decreased compliance of the right ventricle (RV) – or, alternatively, increased stiffness of the RV – during diastolic filling has been shown to correlate strongly with PHT disease progression^6–8^. Clearly, understanding the geometric, structural, and biological factors that contribute to RV stiffening is of significant clinical interest.

The proposed mechanisms driving RV stiffening and impaired diastolic filling range from the molecular to the organ scale. In particular, it has been proposed that RV thickening, extracellular matrix (i.e., microstructural) disorganization, and myocardial remodeling via intracellular and extracellular mechanisms can all contribute to RV stiffening^9–11^. However, the relative contribution of each of these factors to global stiffening of the RV is unclear. Given the predictive power of RV stiffening for PHT outcomes, disentangling the different mechanisms of PHT-induced RV stiffening could reveal novel treatment targets, identify new diagnostic markers, and improve prognosis. However, doing so is not without its challenges. Chief among them is that all the proposed RV stiffening mechanisms occur simultaneously; thereby, obfuscating their individual roles.

Additionally, given the limitations of non-invasive imaging techniques in right heart failure patients, most prior work has relied on non-mechanistic measures of stiffening, such as end-diastolic elastance^4^. In contrast, spatial and directional information could provide additional insight into mechanisms of RV diastolic dysfunction but has largely been ignored. Noteworthy exceptions are recent efforts toward image-based in-vivo RV strain measurements^12^. To overcome the limitations of non-invasive clinical assessment, the development of animal models has allowed for invasive measurements and therefore enabled the study of spatially- and directionally-sensitive properties of the RV^5, 13, 14^. However, most prior animal work has been limited to rodents and other small animal models where the relevance to the human condition is unclear^10, 13, 15–18^.

Thus, the objective of our current work is to use a spatially- and directionally-sensitive repertoire of imaging, experimental, and computational modeling techniques to determine the relative contributions of wall thickening, microstructural disorganization, and myocardial stiffness to RV stiffening. Overall, our goal is to improve our understanding of the mechanisms contributing to PHT-induced RV stiffening with the hope of benefitting clinical practice and right heart failure patient outcomes.

## METHODS

### Animal Procedures

All animal procedures complied with the Principles of Laboratory Animal Care by the National Society for Medical Research. Moreover, all associated protocols abided by the Guide for Care and Use of Laboratory Animals by the National Academy of Science and were approved by the local Institutional Animal Care and Use Committee at Spectrum Health, MI.

In total, our study included twenty male Dorsett sheep (10-12 months of age, Hunter Dorsett Sheep, Lafayette, IN). Eight sheep underwent partial pulmonary artery banding to induce pulmonary hypertension (PHT group, n=8), while the remaining animals served as controls and did not undergo pulmonary artery banding (CTL group, n=12). For pulmonary artery banding, we closely followed the procedure as reported by Verbelen et al^19^. In short, we anesthetized the animals before accessing the pulmonary artery via a left thoracotomy through the 4^th^ intercostal space. Next, we performed epicardial ultrasound to assess biventricular function as well as tricuspid and mitral valvular competence. We then encircled the pulmonary artery with an umbilical tape before its bifurcation. While monitoring systemic and pulmonary pressures via pressure catheters (PA4.5-X6; Konigsberg Instruments, Inc.), we tightened the tape progressively to the brink of hemodynamic instability. After a short observational period, we closed the chest and let the animals recover from the surgery.

After eight weeks of PHT disease progression, we anesthetized the animals, performed a sternotomy and conducted repeated epicardial ultrasound to assess biventricular and heart valve function. Without arresting the heart, we surgically placed a total of 14 sonomicrometry crystals (Sonometrics, London, Canada) across the RV free wall. At the same time, pressure catheters were placed in the right atrium as well as the left and right ventricles. Under open-chest conditions, we recorded sonomicrometry crystal positions as well as atrial and ventricular pressures for at least 10 cardiac cycles. At the end of the procedure, animals were euthanized, and we excised their hearts for downstream imaging and analysis. Control animals underwent the same terminal procedure.

Immediately after excision, we replaced the 14 epicardial sonomicrometry crystals with magnetic resonance imaging (MRI) visible markers (MR-spots, Beekley Corporation, Bristol, CT). The freshly excised hearts were then submerged in phosphate-buffered saline supplemented with the myosin inhibitor 2,3-Butanedione monoxime (Sigma Aldrich) to reduce myocardial contraction^20^. Stored on ice, we shipped the hearts overnight from Michigan to the University of Texas at Austin.

### Magnetic Resonance Imaging

Upon receipt of the excised hearts (approximately 12 hours after euthanasia), we performed detailed MRI scans of each heart. To this end, we first replaced the physiological solution with chilled perfluoropolyether oil (Fomblin 25/6, Solvay Solexis, Thorofare, NJ). Next, we placed the hearts in a 32-channel head/neck receiver coil (Siemens Healthineers, Erlangen, Germany), and scanned them in a Siemens Skyra 3.0 T human MRI scanner using a T1-weighted protocol and a Diffusion Tensor (DT) Imaging protocol; additional details are provided in the supplement.

During postprocessing, we obtained the RV wall anatomy by segmenting the T1-weighted anatomic MRI scans in the open-source software 3D Slicer (Version 4.11). Once segmented, we exported the segmented ventricles to MATLAB (Version R2021) for RV wall volume and thickness calculations. Next, we converted the wall segmentation into an image mask, which we, in turn, imported into the open-source tractography software DSI Studio (June 16 2020 build) for analysis of DT MRI scans; more details are provided in the supplement. After conducting fiber tracking within the RV wall using DSI Studio, we exported the fiber tractography information into MATLAB for additional analysis. There, we subdivided the large RV fiber map into 14 myocardial regions as defined by the epicardial MR-spots. Next, we projected the 3D-fiber orientation vectors within each region onto four equidistant transmural planes between the epicardium and endocardium. Finally, we fit the corresponding projected fiber vectors to a π-periodic von Mises distribution, viz.

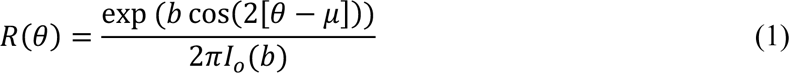

where *θ* is the fiber direction, *b* is the concentration parameter, *μ* is the fiber mean angle, and *I*_0_ is the zeroth order Bessel function. Note that *b=0* represents a random fiber distribution, whereas *bèinf* represents a perfectly aligned fiber distribution.

### Mechanical Testing

Following completion of the MRI protocol, we rinsed the excised hearts and separated the RV wall from the remainder of the heart. Next, we excised myocardial cubes of approximately 10 mm edge size. Each sample was excised such that its edges aligned with the fiber or circumferential direction (F), sheet or longitudinal direction (S), and normal direction (N). Then, we mounted these samples in our custom multi-axial shear tester, which we previously described in detail^20^. Once mounted, we deformed the samples in two simple shear modes up to 30% strain and one tension-compression mode up to 10% strain and recorded the force associated with each deformation. Then we removed the samples from the device, rotated the samples, re-mounted them, and repeated the next two shear modes and tension-compression mode until every unique orientation was tested. Together, our experiments yielded a total of six simple shear and three uniaxial tests per specimen. See Supplementary **Figure S1** for a detailed depiction of each deformation mode.

To quantify the passive stiffness of each specimen, we transformed the force-displacement data into (Cauchy) stress-strain data and quantified stiffness as the slope of the stress-strain curves. To this end, we distinguished between two regimes: i) the low strain regime (i.e., toe stiffness) and ii) the high strain regime (i.e., calf stiffness).

### Histology

After mechanical testing, we fixed samples in 10% neutral-buffered formalin before submitting them to a commercial histology service (Histoserv, Germantown, MD). Histoserv paraffin embedded and sectioned the samples before staining four equidistant, transmural slices with Masson’s Trichrome. Upon receiving the slide images, we identified the relative fraction of collagen by conducting color thresholding as previously described^21, 22^. Thresholding was conducted blinded to avoid bias.

### Quantitative Polymerase Chain Reaction (qPCR)

After mechanical testing, a subset of isolated RV tissue was homogenized using the TissueLyser II system. Total RNA was isolated using the Qiagen RNeasy Mini Kit (Qiagen, 74104) and cDNA was synthesized using the SuperScript IV VILO system (Invitrogen, 11766050). Real-Time qPCR was performed using PowerTrack SYBR Green Master Mix (Invitrogen, A46012) in a QuantStudio 3 Real-Time PCR system. Forward and reverse primers for specific genes of interest were constructed for the sheep genome (*Ovis aries*) and are listed in Supplementary **Table S1**. Relative gene expression was calculated for each sample based on differences in cycle number from the housekeeping genes *GAPDH* and *HPRT1*. Data are presented as normalized relative gene expression, and fold changes between groups were computed from 2^^−ΔCt^ values^23^. Statistical analysis was performed on Δ*Ct* values.

### Finite Element Modeling

We approximated the RV geometry as a quarter sphere with an inner radius of 30 mm, and outer radius of 38 mm, and a maximum applied inner pressure of 40 mmHg. Note that the geometry was later non-dimensionalized, thus, rendering the exact numbers non-consequential. We discretized this geometry with 26,928 linear hexahedral elements using a hybrid formulation as implemented in FEBio (www.febio.org)^24^, see Supplementary **Figure S2** for a mesh convergence study. To model the mechanical behavior of the myocardium, we chose an incompressible Fung-type material model, see more details in the supplement. The parameters for the control case were based on our previous work^20^. The microstructural organization of the control case was based on the same prior work. That is, we assumed that fiber distributions follow a π-periodic von Mises distribution with a dispersion parameter of b=2. Additionally, based on our prior work, we assumed that the mean fiber angle varies transmurally from +22.9° to −76.4° between the epicardium and the endocardium, respectively^20^. We constrained the free surfaces of the quarter sphere to remain within plane; thus, enforcing symmetry conditions. Additionally, we constrained one translational degree of freedom of the node representing the “apex” to avoid rigid body motion. Finally, we simulated the passive filling of the right ventricle under quasi-static conditions using the implicit nonlinear finite element solver FEBio (Version 3.6.0). To study the sensitivity of the diastolic pressure-volume curve (i.e., the filling curve) to wall thickening, microstructural disorganization, and myocardial stiffness we repeated our control simulations after i) increasing the radius of our model by 1.5-fold and 2.0-fold, ii) reducing the concentration parameter *b* to 1.0 and 0.0, see Equation (1), and iii) increasing the stiffness of our constitutive model by 1.5-fold and 2.0-fold.

### Statistics

For statistical comparisons between two independent groups, we first tested for normality via the Shapiro-Wilk test. Where at least one group was not normally distributed, we conducted Mann-Whitney U tests. Otherwise, we conducted independent Student’s t-tests. For comparisons with between and within effects, we used linear mixed models. For comparing mean fiber angles in DT-MRI data, we used a circular mixed model^25, 26^. For all numeric data, we report mean +/− 1 standard deviation when normally distributed or median +/− interquartile range when non-normally distributed. Finally, we used a p-value of 0.05 as our significance level for all inference tests. All descriptive and inference statistical analyses were conducted in MATLAB (Version R2021) except for the linear mixed model analysis, which we conducted in R (Version 2021.09.0).

## RESULTS

### Pulmonary Artery Banding Induced Pulmonary Hypertension in Sheep

Eight weeks of pulmonary artery banding induced significant PHT in sheep as seen in **Table 1**. That is, PHT animals showed increased RV pressure at both end-diastole (ED) and end-systole (ES), as well as increased RV volume (based on ultrasound measurements). The increased RV pressures occurred in the absence of elevated LV pressure or central venous pressure (CVP). Moreover, PHT animals presented with tricuspid regurgitation (TR), reduced RV ejection fraction (EF), reduced fractional area contraction (FAC), and lower tricuspid annular plane systolic excursion (TAPSE); thus, clearly demonstrating reduced RV function.

**Table 1:**
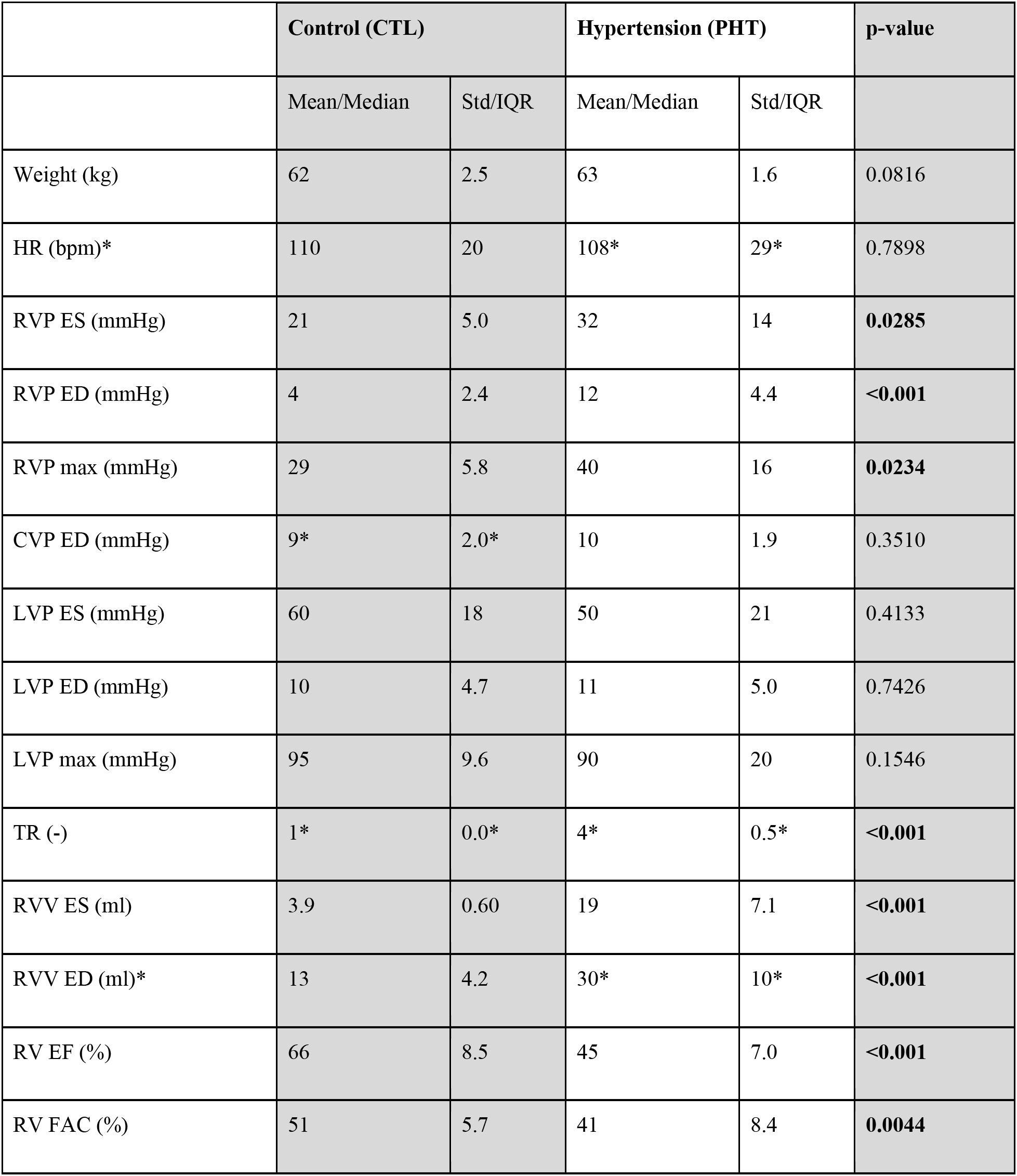
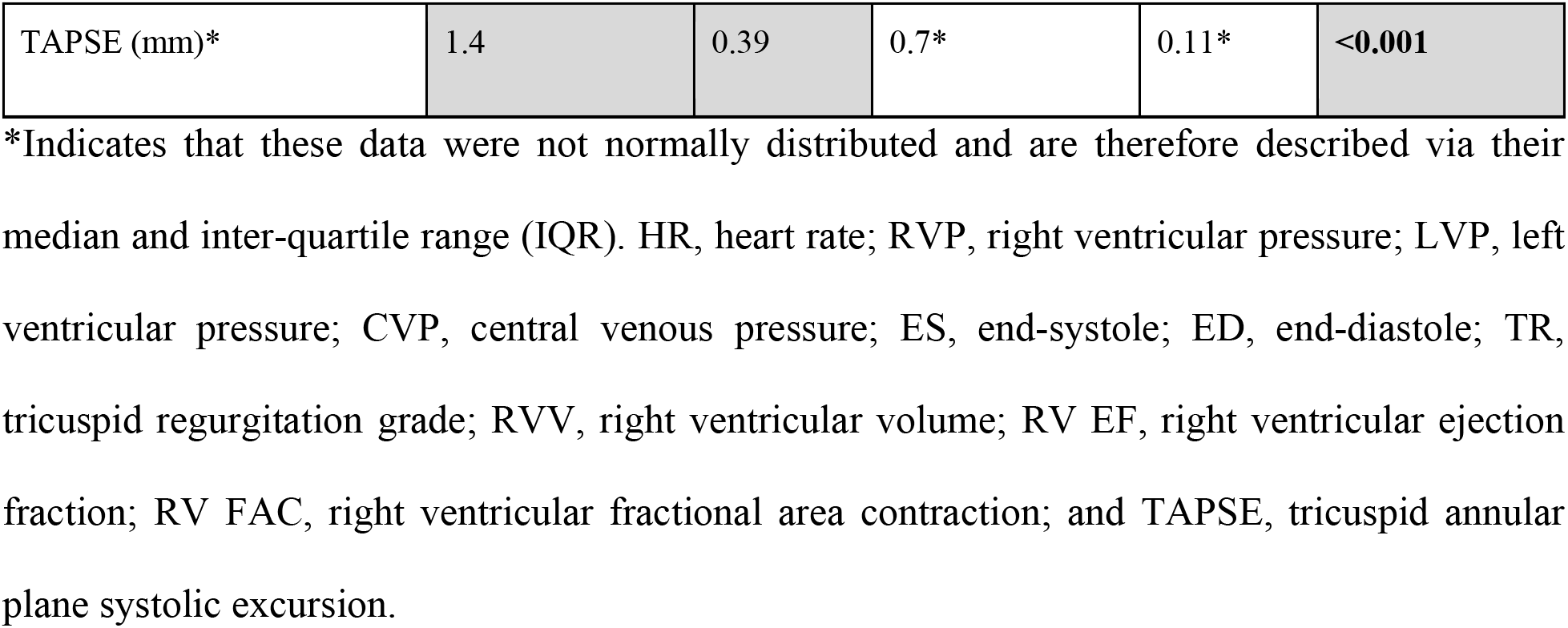
Comparison between measures and right ventricular size and function between the hypertension (PHT) and control (CTL) groups.

|  | Control (CTL) |  | Hypertension (PHT) |  | p-value |
| --- | --- | --- | --- | --- | --- |
|  | Mean/Median | Std/IQR | Mean/Median | Std/IQR |  |
| Weight (kg) | 62 | 2.5 | 63 | 1.6 | 0.0816 |
| HR (bpm)* | 110 | 20 | 108* | 29* | 0.7898 |
| RVP ES (mmHg) | 21 | 5.0 | 32 | 14 | **0.0285** |
| RVP ED (mmHg) | 4 | 2.4 | 12 | 4.4 | **<0.001** |
| RVP max (mmHg) | 29 | 5.8 | 40 | 16 | **0.0234** |
| CVP ED (mmHg) | 9* | 2.0* | 10 | 1.9 | 0.3510 |
| LVP ES (mmHg) | 60 | 18 | 50 | 21 | 0.4133 |
| LVP ED (mmHg) | 10 | 4.7 | 11 | 5.0 | 0.7426 |
| LVP max (mmHg) | 95 | 9.6 | 90 | 20 | 0.1546 |
| TR (-) | 1* | 0.0* | 4* | 0.5* | **<0.001** |
| RVV ES (ml) | 3.9 | 0.60 | 19 | 7.1 | **<0.001** |
| RVV ED (ml)* | 13 | 4.2 | 30* | 10* | **<0.001** |
| RV EF (%) | 66 | 8.5 | 45 | 7.0 | **<0.001** |
| RV FAC (%) | 51 | 5.7 | 41 | 8.4 | **0.0044** |
| TAPSE (mm)* | 1.4 | 0.39 | 0.7* | 0.11* | **<0.001** |

\*Indicates that these data were not normally distributed and are therefore described via their median and inter-quartile range (IQR). HR, heart rate; RVP, right ventricular pressure; LVP, left ventricular pressure; CVP, central venous pressure; ES, end-systole; ED, end-diastole; TR, tricuspid regurgitation grade; RVV, right ventricular volume; RV EF, right ventricular ejection fraction; RV FAC, right ventricular fractional area contraction; and TAPSE, tricuspid annular plane systolic excursion.

### Pulmonary Hypertension Induced Spatially-Dependent Wall Thickening

Using anatomic MRI scans of isolated sheep hearts, we compared both the RV volume and RV wall thickness between PHT and CTL animals. Both metrics were indexed using the animals’ total body weights. **Figure 1A** shows representative segmentations of a PHT (shown in red) and CTL RV (shown in white). We found that PHT induced significant hypertrophy (p=0.001), leading to an average 65% increase in mean RV volume, see **Figure 1B**. Additionally, using our MR spot identification, we found that the RV volume increase was spatially heterogeneous (p<0.0001) and driven by wall thickening primarily at the RV base (markers 1-5; p<0.05) and the RV free wall (markers 9, 12; p<0.01), but not at the apex (markers 7, 11, 14) or the interventricular septum (markers 6, 10, 13), see **Figure 1C**.

**Figure 1:**
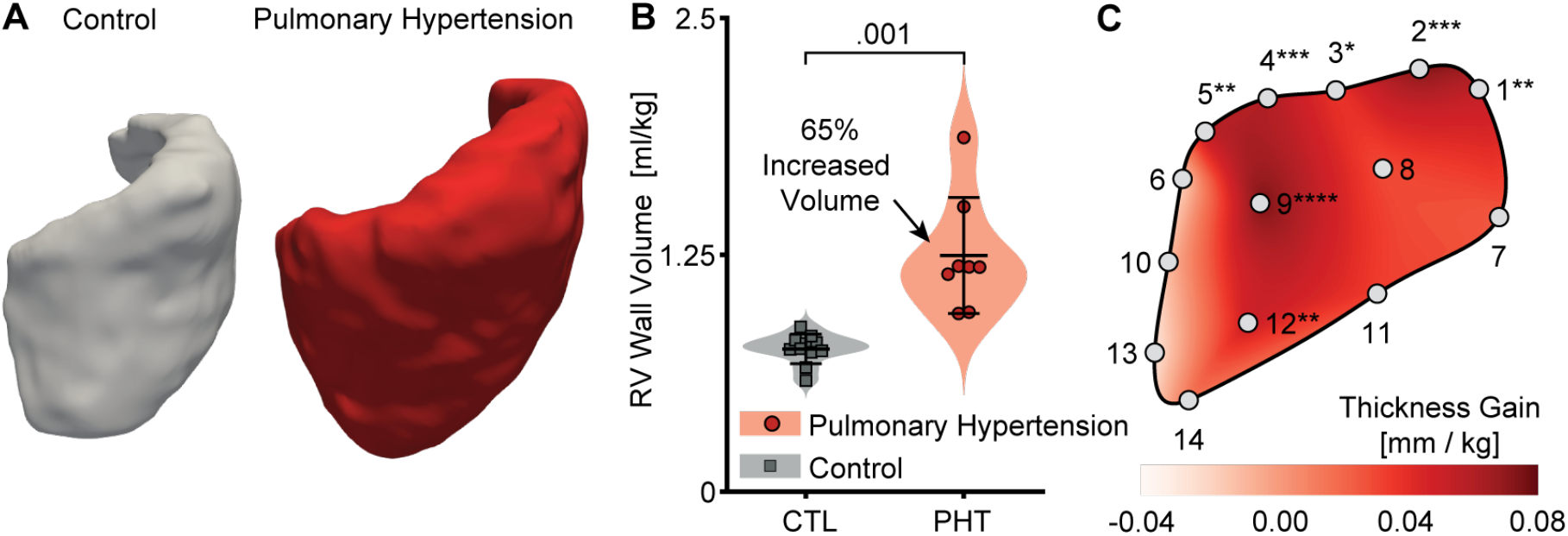
Pulmonary hypertension induced spatially-dependent RV thickening. A) Representative right ventricular segmentations as measured via MRI. B) Pulmonary hypertension induced right ventricular volume increase. C) Spatially-resolved right ventricular thickening computed by comparing the mean thickness map between control (n=11) and pulmonary hypertension (n=8) animals (indexed by animal weight). Base Markers: 1-5, Mid-wall Markers: 8,9,12, Septal Markers: 6,7,10,11,13, Apex Marker: 14. **** (p<0.0001), *** (p<0.001), ** (p<0.01), * (p<0.05)

### Pulmonary Hypertension Increased Extracellular Collagen Deposition

Based on Masson’s Trichrome stains of RV free wall samples, we computed collagen content as area fractions within each field of view. **Figure 2A** shows representative histology images from a PHT animal and a CTL animal demonstrating fibrotic collagen deposition in the former. Note that we included four equidistant transmural sections for each sample in our analysis. We found a significant increase in collagen content (p<0.0001) in each section, see **Figure 2B**. In fact, we found mean increases of 241, 310, 279, and 322% in the epicardial sections (A), mid-wall sections (B) and (C), and the endocardial sections (D), respectively. These histological analyses were further confirmed via transcriptional analysis of hypertrophy-related genes, which revealed a significant increase in *NPPB, MYH7*, and a decrease in *CCN2*, see **Figure 2C**. Furthermore, we conducted a transcriptional analysis of fibrotic gene expression in RV free wall tissue, where *COL1A1*, *COL3A1, MMP2*, and *SDC1* transcription was significantly upregulated in PHT samples, while increases in *FN1* and *TGFB1* are also marginally increased, see **Figure 2D**.

**Figure 2:**
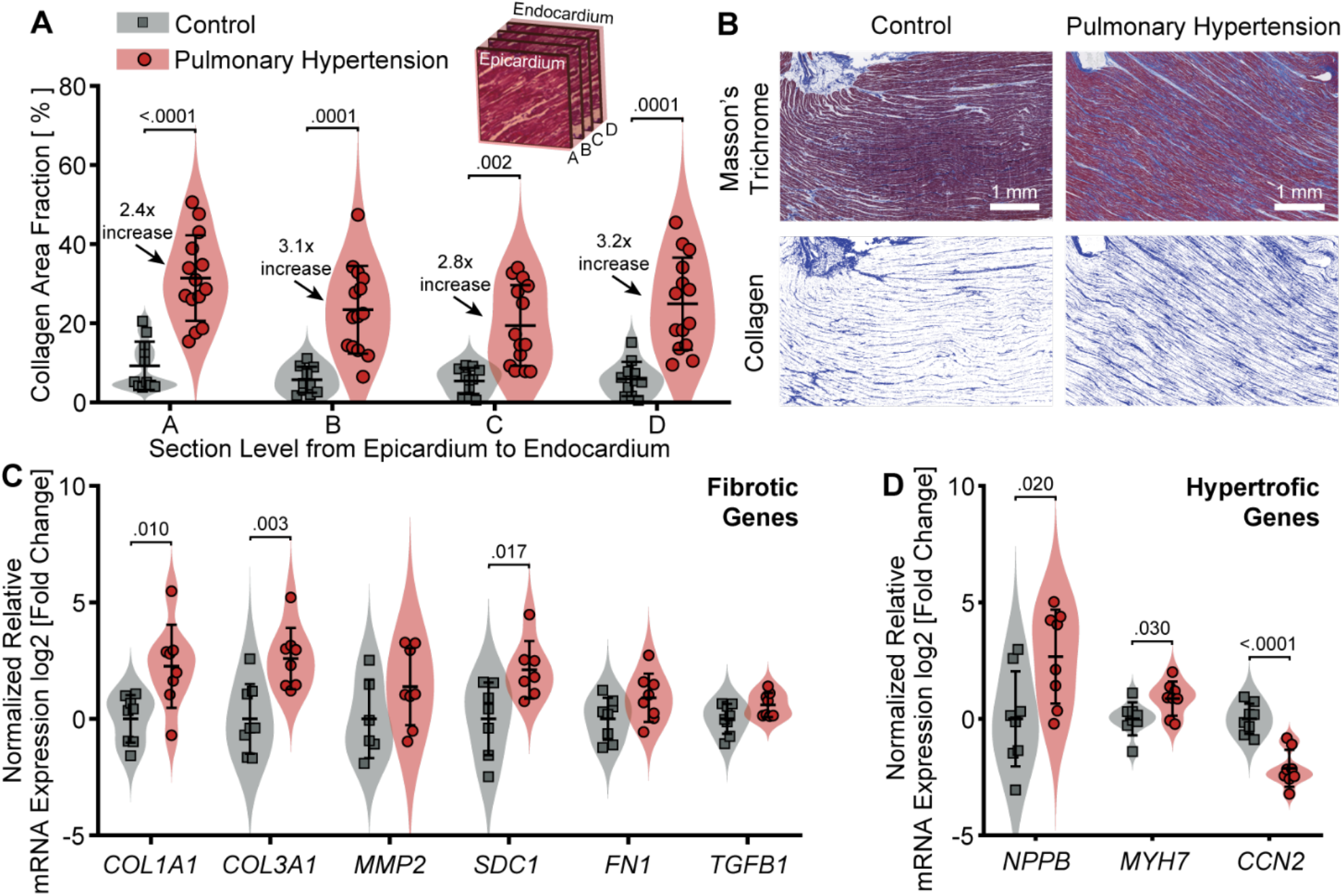
Pulmonary hypertension induced extracellular collagen synthesis. A) Comparison of collagen area fractions in four sections across the myocardial free wall, between control (n=11) and pulmonary hypertension (n=14) specimens. B) Representative Masson’s Trichrome stains of myocardium samples from control and pulmonary hypertension animals. C) Relative expression of fibrosis-related genes. D) Relative expression of hypertrophy-related genes (control n=8, pulmonary hypertension n=8).

### Pulmonary Hypertension Led to Disorganization of Myocardial Microstructure

Using DT-MRI data, we computed the mean myocardial fiber orientation angles and divided the data into four equidistant sections (A-D) across the RV wall (similar to our histological analysis), see **Figure 3A**. In contrast to our histological analysis, we performed the DT-MRI tractography analysis for the entire RV, not just a mid-wall section. By fitting von Mises distributions to the diffusion tensor data, we could capture the mean fiber angle and fiber splay or dispersion. Interestingly, we found that the mean fiber angle did not significantly change between PHT and CTL animals, see **Figure 3B**. However, by comparing the dispersion parameter *b*, see Equation (1), between PHT and CTL animals, we found that myocardial fibers were more dispersed – i.e., less organized – in PHT animals than in CTL animals, see **Figure 3C**. Recall that smaller *b* values imply a larger fiber dispersion.

**Figure 3:**
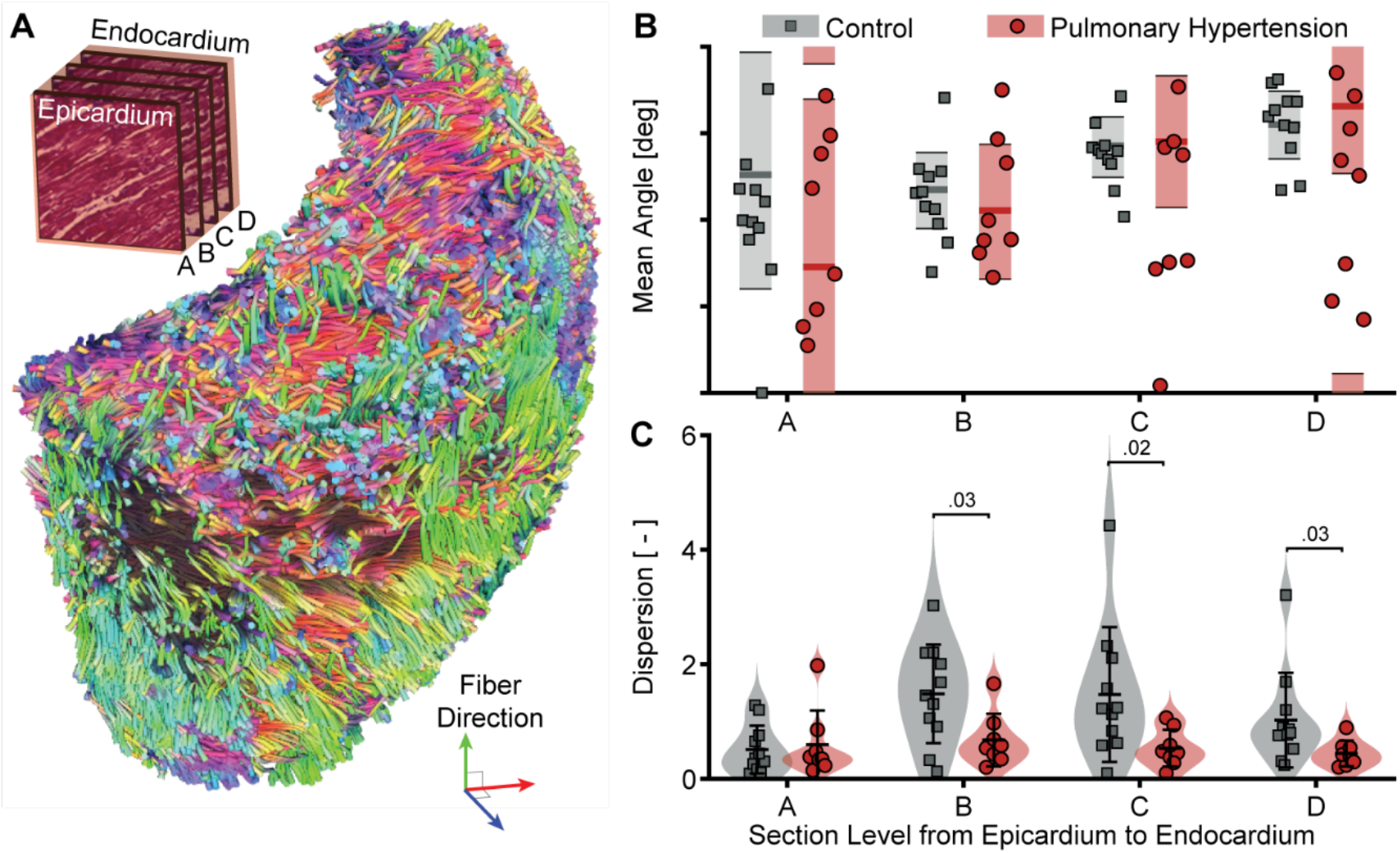
Pulmonary hypertension induced microstructural disorganization. A) Representative diffusion tensor (DT-MRI) tractography with orientation color coding. B) Mean fiber orientation angle in four sections across the right ventricular free wall. Shaded areas indicate the lower and upper bounds of the 95% highest posterior density interval. C) Fiber dispersion in four sections across the right ventricular free wall. (control n=11, pulmonary hypertension n=8).

### Extracellular Collagen Deposition Stiffened Myocardium but is Likely not the Sole Cause

From a comprehensive set of mechanical testing protocols, see **Figure 4A,B**, we found that myocardial stiffness increased in PHT animals relative to CTL. Specifically, we found that the toe-region of samples significantly stiffened under shear, see **Figure 4C,D**, and under extension and compression, see **Figure 4E,F**. That is, myocardium showed stiffening under shear only at small strains (p=0.003), while at large strains no statistical significance was found (p=0.161), see also Supplementary **Figure S3**. In contrast, myocardium showed stiffening under extension and compression both at small strains (p<0.0001) and large strains (p=0.015). Interactions between disease and orientation were not significant. Thus, PHT-induced stiffening was not directionally dependent. Please see Supplementary **Figure S4** for a comparison between the stress-strain curves of control and PHT tissue under all deformation modes. When correlating collagen content with each sample’s stiffness in the fiber direction (FF) we found a statistically significant (p<0.001) and moderately strong relationship (R^2^ = .45), see **Figure 4G,F**. This may imply that increased collagen deposition in the PHT animals is, at least partially, responsible for the increased stiffness of the tissue; but not solely.

**Figure 4:**
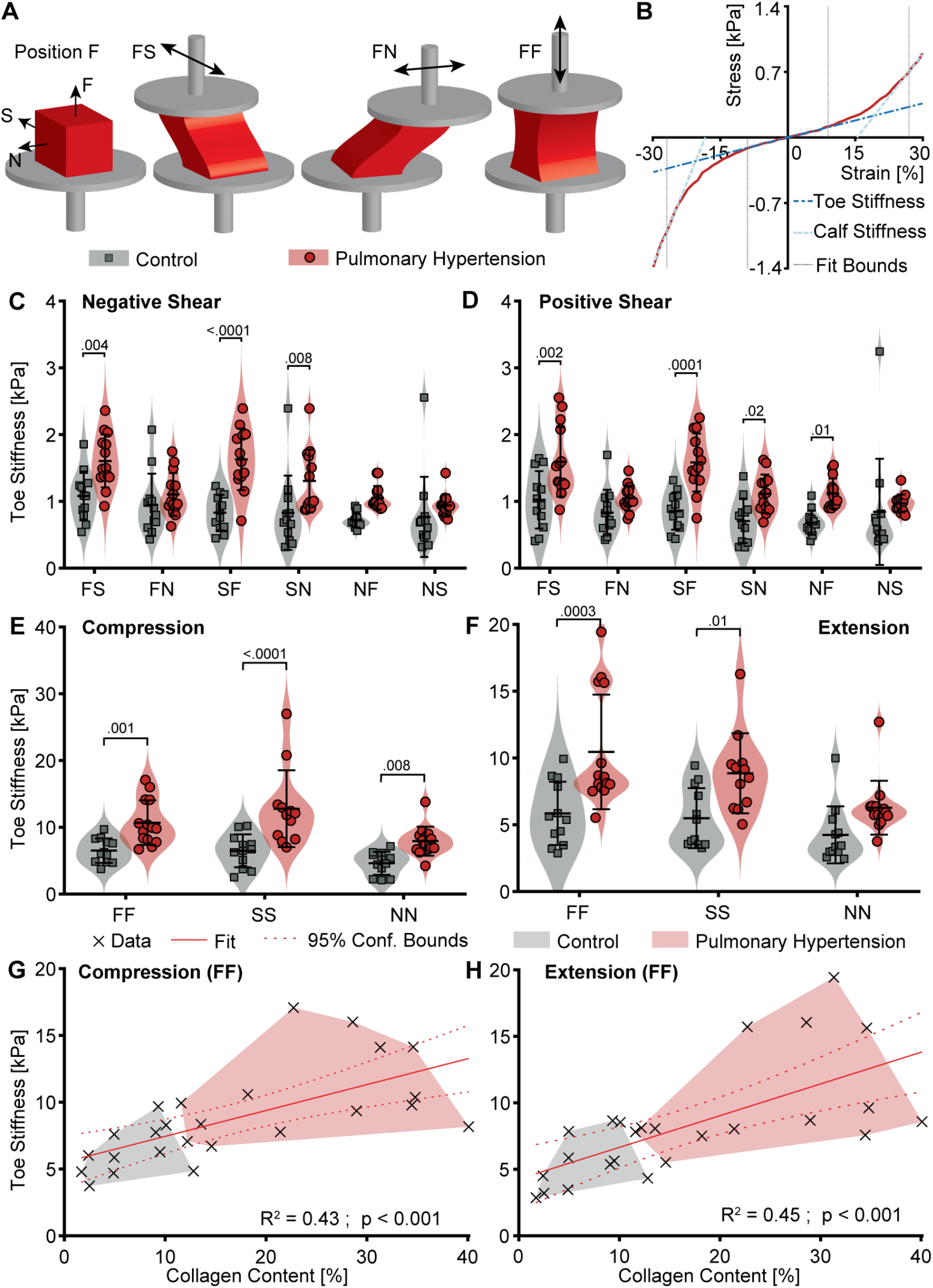
Pulmonary hypertension induced myocardial stiffening. A) Nomenclature for mechanical testing modes. B) Representative stress-strain data. C-D) Toe stiffness (i.e., stiffness at small strains) in six shear directions in positive and negative direction. E-F) Toe stiffness under compression and tension in three material directions. (control n=12, pulmonary hypertension n=14). G-H) Correlation between fiber compressive and tensile stiffness in fiber direction (FF) with collagen content (per area fractions) (control n=11, pulmonary hypertension n=14).

### Myocardial Stiffening is a Significant if not Primary Cause for RV Stiffening

Our data revealed three potential mechanisms of RV stiffening: i) thickened RV walls, ii) a disorganized microstructure, and iii) stiffened myocardium. To decouple these three mechanisms, we built finite element models of a simplified RV and used the models to simulate end-diastolic filling, see **Figure 5A**. First, we only increased the wall thickness of our model without changing any other parameters. Even at an increase of 50% and 100%, we found only relatively small changes to the passive filling curve, see **Figure 5B**. In fact, at a normalized pressure of 2.5 (∼40mmHg) a 2-fold increase in wall thickness reduced the total filling volume (*V_f_*) only by 2.8%. Similarly, we drastically altered the myocardial organization and increased its fiber dispersion while keeping all other parameters the same, see **Figure 5C**. Again, we found only small effects on the passive filling curve. That is, at a normalized pressure of 2.5 (∼40mmHg), partial, and even full dispersion of the collagen matrix lead to a modest reduction of filling volume of only 2.2%. Finally, we also increased the myocardial stiffness, see **Figure 5D**. This change significantly altered the filling curve reducing *V_f_* by 11.7%; thus, significantly exceeding the effects of wall thickening and microstructural disorganization.

**Figure 5:**
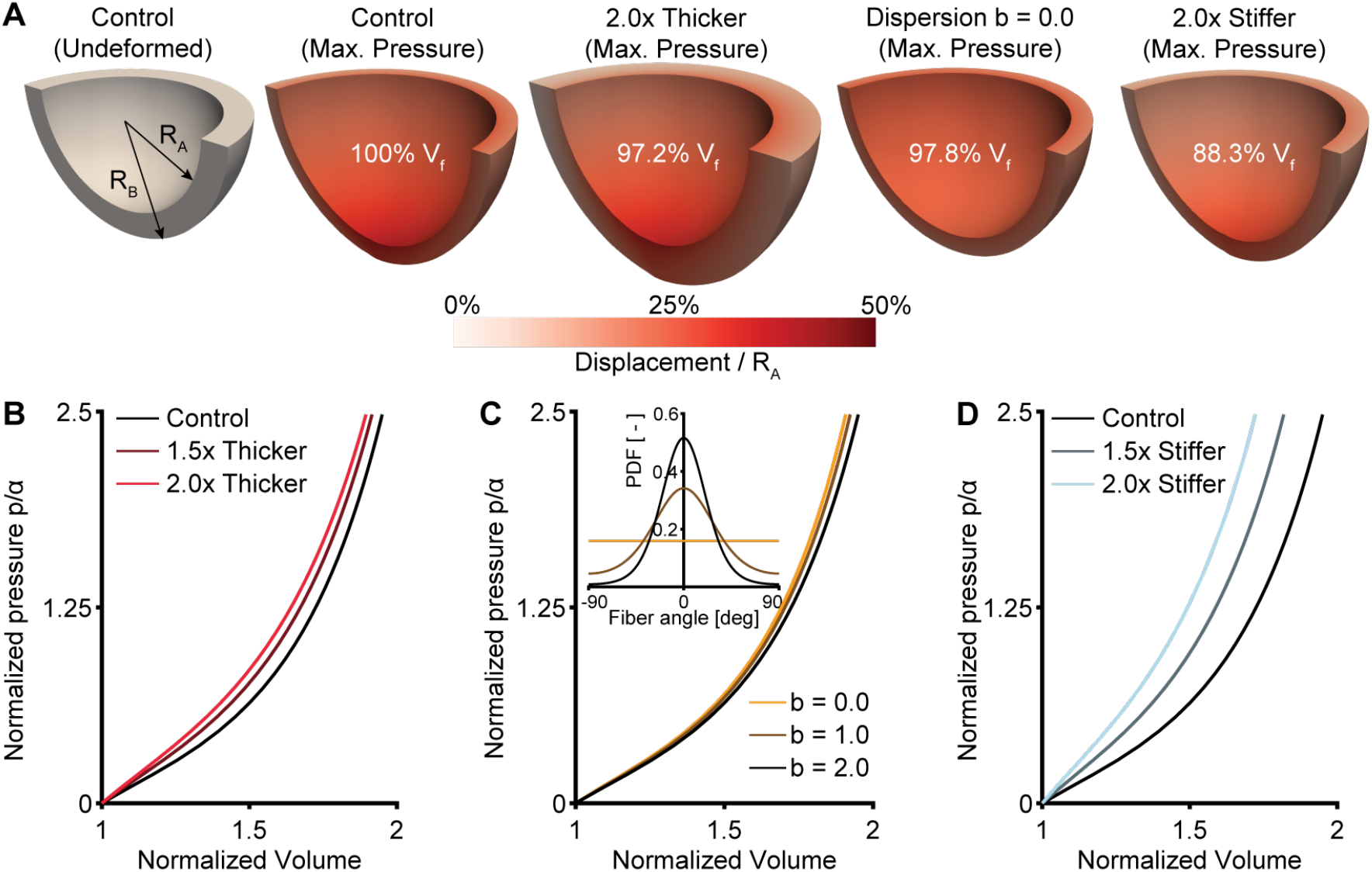
Myocardial stiffness is the primary wall stiffening mechanism as predicted by a computational model of the right ventricle. A) Simplified model of the right ventricle under zero pressure (grey) and at max pressure (red). R_A_ and R_B_ denote the inner and outer wall radius, respectively. We compared the diastolic filling volume (*V_f_*) after thickening the wall by up to 2.0-fold, increasing the fiber dispersion until random (b=0.0), and increasing myocardial stiffness by up to 2.0-fold to the filling volume of the control ventricle. B-D) Passive filling curves of the right ventricle after wall thickening, changing fiber dispersion, and increasing myocardial stiffness, respectively. Pressures (*p*) are normalized by the initial myocardial stiffness (*a*), while volumes are normalized by the pressure-free volume.

## DISCUSSION

Our goal for this work was to use spatially- and directionally-sensitive measures of RV remodeling in a large animal model of PHT to delineate the relative importance of the different RV stiffening mechanisms of wall thickening, fiber orientation, and myocardial stiffness. This work provided new insights into the mechanisms of RV stiffening.

We confirmed that the RV adapts to hypertension via significant volumetric growth. Interestingly, this volumetric growth is not uniform. Instead, we found a markedly heterogeneous thickening. Specifically, the base and the free wall of the RV were thickened in the PHT animals, while the apex (e.g., markers 13 & 14 in **Figure 1C**) and the wall near the interventricular septum (e.g., markers 6, 7, 10 & 11 in **Figure 1C**) were nearly unchanged. Likely, this uneven distribution stems from differing growth stimuli across the wall. At the apex and the septum, where the RV is tied closely to the LV, mechanobiological stimuli are likely smaller than in the untied free wall and at the base. This hypothesis agrees well with the general understanding that it is the lateral RV wall that dilates in RV failure and is implicated in tricuspid valve regurgitation^27^. Of note, others have made the opposite argument. Specifically, Andersen et al. concluded their meta-analysis on RV fibrosis with the observation that late gadolinium enhancement – as a measure of myocardial fibrosis – was found primarily near the RV insertion point at the interventricular septum and argued that higher local stresses may be the cause for those local hotspots^28^. Our contrasting findings imply that local hypertrophic thickening may not necessarily correlate with local fibrosis and that both phenomena should be viewed as separate stiffening mechanisms.

Regarding RV fibrosis, we confirmed that PHT induces significant collagen synthesis. Notably, we found an increase in interstitial collagen of more than 60% in PHT tissue relative to CTL. This histological finding was consistent throughout the RV wall thickness and showed signs of persisting as several genes encoding fibrotic proteins continued to be upregulated in the myocardium at 8 weeks after pulmonary artery banding. This contrasts findings in humans where interstitial RV collagen accumulation has been described as minor^6, 29^. One possible explanation for this difference in RV fibrosis across species is the fast onset of PHT in our animal model as opposed to the gradual onset of disease in (some) human conditions. It is possible that the fast adaptive response of the sheep RV may result in aberrant collagen synthesis as opposed to the more chronic natural history of human disease. This argument is supported by the findings of Rain et al., who observed that in rats with mild dysfunction, the RV stiffened primarily via hypertrophic thickening. However, in rats with severe RV dysfunction the ventricle stiffened primarily via myocardial stiffening as a result of collagen deposition^10^. Note that we evaluated collagen density as a measure of RV fibrosis only on mid-wall samples and thus cannot compare our findings to the reports based on late gadolinium enhancement-based spatial maps of fibrosis, as reported above.

When characterizing microstructural organization, we found that mean fiber directions across the RV had not changed, but that fiber dispersion had increased. We submit that increased fiber dispersion is likely a result of newly synthesized collagen being deposited into a pre-existing, pathologically deformed extracellular network. However, these findings directly contrast with others’ prior studies in rodents that observed the opposite behavior – mean fiber direction was changed and fiber alignment increased^13, 14^. The source of this disagreement is unknown and may be attributed to the difference between the natural history of RV remodeling in rodents and in our large animal model. That said, the current findings agree well with our prior work where we found tricuspid valve leaflets underwent the same fibrotic changes in animals with RV failure^30^ and are consistent with a general understanding of microstructural remodeling associated with fibrosis^31^.

Importantly, our large animal model of PHT confirmed several previous findings that pulmonary hypertension induces myocardial stiffening^10, 13, 32, 33^. Though, the current study is the first to subject maladapted RV myocardium to multi-directional shear loading scenarios. We found that RV myocardium had significantly stiffened in almost all shear directions. Additionally, myocardium stiffened under compression and tension in (nearly) all directions. At the same time, the measured stiffness increase in the fiber direction correlated significantly with collagen content. Finally, we found that stiffness significantly differed only at small strains (i.e., toe stiffness), but not at large strains (i.e., calf stiffness). Recall, there are two broad contributors to myocardial stiffness: collagen-dominated extracellular stiffness and titin-dominated cellular stiffness^6^. The former depends on collagen type, cross-link density, collagen density, and collagen alignment^34–36^. The latter depends on titin isoform ratio and the state of titin phosphorylation^37^. Titin is activated even at small strains, while collagen is activated only at moderate to large strains^5, 13^. Our observation that stiffness correlated moderately with collagen content implicates collagen as a significant contributor, but not the sole driver of myocardial stiffening. In addition, the dominance of small strain stiffening implicates titin as another possible important driver of stiffness. The absence of cell-level stiffness measurements in the current study, however, makes it difficult to draw detailed conclusions regarding the extent of titin-dependent intracellular myocardial stiffening mechanisms in our model of PHT.

Finally, we conducted a finite element analysis of RV diastolic filling and used this computational model to independently investigate the impact of each proposed stiffening mechanism – wall thickening, microstructural disorganization, and myocardial stiffening – on RV diastolic stiffness. Interestingly, we found that of the three mechanisms, myocardial stiffening had the largest influence. To ensure that our findings weren’t the result of poor modeling assumptions, we conducted sensitivity analyses and varied all elements of our model. That is, we tested different geometries, material models, and assumptions of the organization of the myocardium model. None of these numerical permutations fundamentally altered our findings. In support of our findings, Baicu et al. and Kwan et al. suggested that hypertrophic thickening is primarily an early adaptive mechanism that improves systolic function, while myocardial stiffening is a late adaptive mechanism that improves the RV’s resistance to excessive dilation^18, 33^; hence, concurring with our observations that myocardial stiffness is the primary mechanism driving RV stiffening.

### Clinical Significance

The clinical significance of our work stems from several key points. First and foremost, we have demonstrated the immense remodeling potential of the RV in a large animal model. That is, after only 8 weeks of PHT the RV grew on average by 65% in volume, remodeled its microstructure, and fundamentally altered its intrinsic material properties. Second, we showed that RV hypertrophy is driven primarily by growth at the base and free wall, while all other areas seem to be protected from excessive growth by the left ventricle. From a clinical perspective, this suggests RV hypertrophy should be evaluated with spatial heterogeneity in mind. That is, hypertrophy should be evaluated where the ventricle is likely most remodeled. Most importantly, our findings suggest that the inherent stiffness of the RV myocardium may be the primary driver of RV stiffening. Thus, myocardial stiffening via intracellular (titin activation) or extracellular mechanisms (collagen deposition) should be considered a prime candidate for predicting RV diastolic dysfunction. While there are global measures of RV stiffening, such as end-diastolic elastance, other direct and spatially-sensitive methods such as ultrasound- or MRI-based strain measurement techniques should be explored for the RV myocardium^38^. Finally, our work suggests that collagen synthesis and extracellular matrix fibrosis are critical drivers of myocardial stiffening of the RV. This is of potential clinical significance as modern MRI techniques are sensitive to collagen signatures and could be further used to non-invasively estimate material stiffness of the RV and to predict clinical outcomes of pulmonary hypertension and heart failure^39^.

### Limitations

The primary limitation of our study is that the onset of PHT in our animal model is sudden rather than progressive as in most clinical instances^40^. For example, others have found in rodents some evidence that suggests that sudden onset PHT may have different pathology than progressive onset PHT^18, 33^. Additionally, we have captured only one time point at eight weeks of disease progression. Thereby, we are unable to characterize the temporal evolution of RV stiffening in this large animal model. We also did not conduct cell-level studies that could have provided further insight into the intracellular mechanisms of myocardial stiffening and potentially explain the relative importance of extracellular and intracellular contributions. Finally, our study was conducted on sheep and may not fully resemble the human clinical condition.

## Novelty & Significance

What is known?

- Pulmonary hypertension leads to hypertrophy and diastolic stiffening.
- Diastolic stiffening is a predictor of disease progression.
- Multiple mechanisms on multiple scales contribute to diastolic stiffening.

What new information does this article contribute?

- Mechanisms of diastolic stiffening are spatially- and directionally-dependent.
- Myocardial stiffening, i.e., stiffening of the myocardium itself, is likely the major driver of diastolic stiffening.

We found that PHT induces RV wall thickening, microstructural disorganization, and myocardial stiffening. These mechanisms were both spatially and directionally dependent. Most critically, we observed – through experimental measures of fibrosis and numerical simulations of diastolic filling – that myocardial stiffening is the most important contributor to RV stiffening. Thus, by extension, myocardial stiffening may be an, if not the most, important predictor for PHT progression. Given the significant correlation between myocardial stiffening and collagen synthesis, extracellular matrix-sensitive imaging modalities may be used as a future clinical proxy for myocardial stiffness and a useful tool for evaluating disease outcomes.

## ACKNOWLEDGEMENT

MKR acknowledges the partial support from the National Institutes of Health via grants R01HL165251 and R21HL161832, and the National Science Foundation via grant 2127925. TAT acknowledges the partial support from the National Institutes of Health via grants R01HL165251 and R21HL161832. MRB acknowledges support from the National Institutes of Health via grant R00HL146951. MM acknowledges support from the American Heart Association via its predoctoral fellowship 902502.

